# Development and Use of miRNA-derived SSR Markers to Study Genetic Diversity, Population Structure and Characterization of Genotypes for Heat Tolerance Breeding in Bread Wheat (*Triticum aestivum* L.)

**DOI:** 10.1101/2020.03.17.995142

**Authors:** Sandhya Tyagi, Anuj Kumar, Tinku Gautam, Renu Pandey, Reyazul Rouf Mir

## Abstract

Wheat is one of the most important cereal crop in the world. Heat stress is an important abiotic stress limiting wheat production and productivity in the world including south-east Asia. The importance of miRNAs in gene expression under various biotic and abiotic stresses is well documented. Molecular markers, especially SSR markers, plays an important role for the success in molecular plant breeding programs. The discovery of SSRs from non-coding regions has been a challenging task. Therefore, development of novel miRNA-based SSRs from the conserved portions of the genome will prove useful for the study of genetic diversity of heat-responsive miRNA-genes in wheat. In the present study, efforts are made to mine SSR markers from 96 members of heat-responsive miRNA-genes of wheat followed by their validation using 37 contrasting (heat tolerance/susceptible) wheat genotypes. Among a set of 13 miRNA-SSRs used,7 SSRs were found polymorphic. Among these polymorphic SSR markers, three found to be very informative SSRs (HT-169j, HT-160a and HT-160b) and could largely discriminate heat tolerant genotypes from the heat susceptible ones. Further analysis based on Polymorphism Information Content (PIC) revealed that miRNA genes were more diverse in susceptible genotypes compared to tolerant genotypes. Ours is the first report that the genic/miRNA markers could be successfully used to study wheat diversity, population structure and characterization of trait specific germplasm. The important and useful miRNA-based SSRs, therefore, would serve as best markers in the marker-assisted breeding programs aimed at enhancing heat tolerance of Indian wheat.

## Introduction

During the past two decades, a variety of ‘Omics’ tools and techniques have been deployed in crop improvement programs. RNA interference (RNAi) technology is considered one of the most important tools for genetic manipulation/ functional genomics. The phenomenon of RNAi involves cleavage of double-stranded RNA (dsRNA) [1] that produces two small non-coding RNAs known as small interfering RNA (siRNA) and microRNA (miRNA). MicroRNAs (miRNAs) play a key role in gene expression alteration under various biotic and abiotic stresses. In addition, miRNAs play a major role in growth, development, and response to various biotic and abiotic stresses. The pace of discovery of small miRNAs have increased over the years due to the advances made in the area of next-generation high-throughput sequencing technologies that open up possibilities of exploring small RNA (sRNA) populations in economically important crop species like wheat [2,3].

Heat stress is considered as one of the most significant abiotic stresses that negatively impact yield and yield components in crop plants [4]. Wheat, a staple cereal crop in the world, is highly sensitive to heat stress. The production and productivity of wheat is greatly hampered by heat stress. Therefore, there is an urgent need for exploration and utilization of novel genetic resources to improve tolerance/sensitivity level in cultivated bread wheat genotypes for mitigating the damaging effects of heat stress [5]. Efforts are also needed to explore the possibility of using small RNA molecules like miRNAs and miRNA derived molecular markers including miRNA derived SSR markers for breeding heat tolerant wheat varieties.

Several QTL studies have been conducted in the past for the exploitation of the genetic variation among the existing cultivars for stress tolerance in different crops [7,5,9]. Molecular markers, especially co-dominant markers like simple sequence repeat (SSRs), flanking these reported QTLs play an key role in marker-assisted wheat breeding programs. SSRs are important and widely distributed in plant genome, especially in coding and non-coding genomic regions [10]. Most of the earlier reported SSRs are from the protein-coding regions of the wheat genome [11]. As per published data, SSRs from non-coding regions of the wheat genome have not been fully exploited, catching attention of a scientific community towards thediscovery of SSR markers from non-coding regions.

Genome-wide analysis of miRNAs for different traits has been conducted in several plant species, including rice, maize, barley, and Arabidopsis [12,13,14,15,16,7]. In wheat, the genomewide analysis of miRNAs has also been reported under heat stress condition [17,18,19]. However, to the best of our knowledge there is no report available on the development and use of markers based on miRNAs (miRNA-SSRs) in wheat. Therefore, in this study, an attempt was made for genome-wide identification and characterization of miRNA-SSRs for heat tolerance in wheat. We developed SSR markers from the conserved microRNA (miRNA) genes in wheat for heat sensitivity to characterize a diverse set of wheat genotypes. These miRNA gene based SSRs (miRNA-SSRs) can be used in marker aided selection (MAS) programme for heat tolerance in wheat. Further, evolutionarily miRNAs are highly conserved and hence, miRNA-SSRs can reveal more diversity compared to existing markers among closely related genotypes or self-pollinated crop species.

## Materials and Methods

### Plant materials and DNA isolation

Thirty-seven (37) diverse wheat genotypes comprising both heat tolerant (26) and heat susceptible (11) genotype were used in this study (Table 1). The heat tolerant and susceptible genotypes are being used as national donors and recipients in development of different genetic resources including mapping populations and in discovery of QTLs/genes for heat tolerance and related traits. DNA isolation was carried out following the procedure described by Saghai-Maroof et al. [20]. After RNAase-treatment, DNA quality and purity were checked on 0.8% agarose 1× TAE gel. DNA quantification was done using Thermo Scientific NanoDrop 1000 spectrophotometer and diluted to the final concentration of 25 ng/μl.

**Table 1:** Details of 37 heat tolerant/susceptible wheat genotypes used in the study.

| S. No | Genotype | Tolerant vs Susceptible | Pedigree | Name of developing centre | Area/zone of Adoption |
| --- | --- | --- | --- | --- | --- |
| 1 | Lok-1 | Tolerant | S-308 x S-311 | SANOSARA | CZ |
| 2 | HUW-234 | Tolerant | HUW 12*2 / CPAN 1666//HUW12 | - | NEPZ |
| 3 | VL-946 | Tolerant | HW 2045/VW 2185 | IIWBR, Karnal | - |
| 4 | JO-07-47 | Tolerant | HEAT TOLERANT NURSERY 2015-16 | IIWBR, Karnal | - |
| 5 | HD-3043 | Tolerant | PJN/BOW//OPATA*2/3CROC_1/A. Squarrosa(224) //OPATA | ICAR-IARI, New Delhi | NWPZ-TS, RIR |
| 6 | GW2008-153 | Tolerant | HEAT TOLERANT NURSERY 2015-16 | IIWBR, Karnal | - |
| 7 | NIAW-2477 | Tolerant | HD2402/VL639 | - | NWPZ |
| 8 | PBW-51 | Tolerant | PBW 51*3/ KS90 H 450/ Moro // HW4444 (HUW234*3/CLr19) | - | - |
| 9 | RAJ-4037 | Tolerant | DL788-2/RAJ3717 | RAU, Durgapur | IR, TS |
| 10 | BARKARA | Tolerant | Viginta/ Danubia | - | - |
| 11 | BWL-0924 | Tolerant | N/A | - | - |
| 12 | SONARA | Tolerant | YT54/N10B//2*Y54 | New Delhi | NPZ/CZ |
| 13 | GIZA-168 | Tolerant | Mil/Buc//Seri | Egypt | - |
| 14 | BWL-1771 | Tolerant | N/A | - | - |
| 15 | TEPORO | Tolerant | - | - | - |
| 16 | HD-2864 | Tolerant | DL 509-2/DL 377-8 | New Delhi | CZ |
| 17 | CHIRYA-3 | Tolerant | - | - | - |
| 18 | WH-730 | Tolerant | CPAN 2092/Improved Lok I | CCSU, Hisar | - |
| 19 | HI-1563 | Tolerant | MACS2496*2/MC10 | - | - |
| 20 | DBW-14 | Tolerant | RAJ 3765/PBW 343 | DWR | NEPZ |
| 21 | RAJ-4083 | Tolerant | PBW343/UP2442/WR258/UP2425 | SKNAU-RARI, Durgapura | PZ-LS,IR |
| 22 | IC-32586 | Tolerant | - | - | - |
| 23 | IC-118737 | Tolerant | - | - | - |
| 24 | BWL-1793 | Tolerant | ND/VG9144 | - | Germplasm line |
| 25 | RAJ-3765 | Tolerant | HD 2402/VL 639 | RAU Durgapur | NEPZ |
| 26 | HD-2967 | Tolerant | ALD/COC//URES/HD 2160 M/HD 2278 | ICAR-IARI, New Delhi | - |
| 27 | HUW-510 | Susceptible | HD 2278 / HUW 234 // DL 230 - 16 | - | PZ |
| 28 | RAJ-4014 | Susceptible | DL 8025/K 9011 | Rajasthan | - |
| 29 | HUW-468 | Susceptible | CPAN 1962 / TONI // LIRA / PRL | - | NEPZ |
| 30 | HD-2824 | Susceptible | - | - | NEPZ |
| 31 | PBW-343 | Susceptible | PBW 343*3/ KS 90 H 450/Moro// HW4444(HUW234*3/CLr19) | - | NWPZ |
| 32 | Raj- 3184 | Susceptible | - | - | - |
| 33 | DL-784-3 | Susceptible | KAL*4/TR380.27*4/3AG/3/HD2281 | IARI, New Delhi | NEPZ |
| 34 | HD-2402 | Susceptible | HD2177//CN067/BB/3/HD 2160/4/HD 2236 | - | NEPZ |
| 35 | GW-322 | Susceptible | GW173/GW196 | JAU, Junagarh | IR, TS |
| 36 | HD-2327 | Susceptible | - | - | CZ |
| 37 | HS-240 | Susceptible | AU /KAL- BB//WOP'S'/P AVON'S' | - | NHZ |

### Heat-responsive miRNA-SSRs mining and validation

Extensive literature survey was conducted to identify heat-responsive miRNAs from cereal crops. Reference miRNAs were downloaded from miRBase21.0 (http://www.mirbase.org/) [21] and fulllength genomic transcripts representing pre-miRNA were obtained in FASTA format using BioMart-Ensembl genomes mining tool [22] available on Ensembl Plants platform [23]. In order to identify the 500 bp upstream and 500 bp downstream of the premature-miRNA sequence (1000 bp long primary miRNA sequence), BLASTn algorithm were conducted against latest release of wheat genomic data available on Ensembl Plants. BLAST algorithm was conducted as earlier described by Mondal et al. 2014 (16) and Kumar et al. 2017 (24); i) premature miRNA sequences of wheat were directly used as ‘query’, ii) in case of other plants, corresponding mature miRNAs were taken to find the wheat miRNA orthologue and then respective premature sequences of rice miRNA were used as ‘query’, iii) premature miRNA sequence of other plants were used directly as a query in case that mature miRNA orthologue are absent to wheat miRNA database. As a result of BLAST search, different wheat primary miRNA sequences of 1000 bp length were selected and further scanned for mining the repeats using Simple Sequence Repeats Identification tool (SSRIT) [25] (http://archive.gramene.org/db/markers/ssrtoolmarkers/ssrtool) with default parameters.

After that, the extended 500 upstream and 500 downstream sequences of miRNAs with a repeat motif more than N6 were chosen for designing SSR primers using online software program PRIMER3.

### SSR marker genotyping and data recording

PCR amplification of a set of 37 genomic DNAs using different identified miRNA-based SSR primers was carried out using a reaction mixture of 20 μl containing 50 ng template DNA, 0.25 μM of each primer, 0.02 mM dNTPs, 1× PCR buffer (10 mM Tris–HCl pH 8.4, 50 mM KCl, 1.8 mM MgCl_2_ and 0.01 mg/ml gelatine) and 0.5 U Taq DNA polymerase. The PCR reaction was performed using a C1000 Touch™ Thermal Cycler (Bio-Rad Laboratories, Hercules, CA, USA) with the following profile: an initial denaturation step of 95°C for 5 min followed by 34 cycles of 95°C for 45seconds, annealing for 30 seconds, and 72°C for 45 seconds. The final extension was at 72°C for 7 min. Amplified products were resolved on 10% PAGE using standard procedure. The amplified products were resolved on 10% poly-acrylamide gels using vertical gel electrophoresis unit of Bangalore Genei (glass plate size 19 × 17.5 cm). After electrophoresis, the gels were stained using silver staining [26]. For each SSR, the data was recorded on the basis of an allelic fragment size in two different sets of genotypes. Absence of a band was recorded as a null allele.

### Analysis of population structure

A model-based (Bayesian) method with 13 miRNA-SSRS was utilized to evaluate the possible number of sub-populations in the population of 37 candidate wheat genotypes/lines. The population structure analysis was performed using software program STRUCTURE 2.3.4 [27]. Population structure was analysed by setting the number of sub-populations (k-values) from 1 to 10 and each run was repeated five times. The program was set on 100,000 as burn-in iteration, followed by 150,000 Markov chain Monte Carlo (MCMC) replications after burn in along with admixture model [28]. The STRUCTURE HARVESTER web version v0.6.94 [29] was used to work-out the exact number of sub-populations using modified delta K (ΔK) method, which provides real number of clusters [30]. Within a group, genotypes with affiliation probabilities (inferred ancestry) ≥ 80% were assigned to a distinct group, and those with < 80% were treated as admixture, i.e. these genotypes seem to have a mixed ancestry from parents belonging to different gene pools or geographical origin.

### Diversity analyses and population differentiation

To demonstrate the use of miRNA based SSR markers for the study of genetic diversity, a set of 13 miRNA-SSR loci was used to generate genotypic data on 37 wheat genotypes. For this purpose, genotypic data was statistically analysed using software program POPGENE version1.31 [31] to measure the total number of alleles (Na), the number of effective alleles (Ne) and Shannon’s Information index. We calculated the Polymorphic Information Content (PIC) for each SSR marker loci using the software package Power-Marker version 3.25 [32]. Dissimilarity matrix was calculated with DARwin version 6.0 (DARwin: Dissimilarity Analysis and Representation for WINdows; [33] using genotypic (allelic) data. Calculated dissimilarity matrix was used for clustering of 37 genotypes and dendrogram preparation. An un-weighted neighbour-joining (UNJ) method [34] was followed for this by bootstrap analysis with 1000 permutations. The genotypic data (multiple loci and multiple genotypes) was also analysed using GenAlEx version 6.5 software [35] for conducting principal coordinate analysis (PCA). A pair-wise genetic distance matrix was calculated and used for preparing PCA plots. Other genetic diversity parameters such as pair-wise Nei’s unbiased genetic distance and genetic identity [36], and gene flow (Nm) were also computed using software GenAlEx version 6.5.

### Analysis of molecular variance (AMOVA) and genetic diversity indices

Analysis of molecular variance (AMOVA) was performed to calculate the level of genetic variation not only among the genotypes but also within the populations. The genetic distance matrix (same as used for PCA) was utilized for conducting AMOVA. The number of subpopulations determined based on STRUCTURE results were used for AMOVA. The AMOVA was performed using software GenAlEx6.5 [35].

## Results

### Discovery and validation of heat-responsive miRNA-SSRs in wheat

Based on the literature search, we identified 39 miRNA family comprising 96 members responsive to heat stress in various crops including Arabidopsis, rice and wheat. Out of 39 families, only 14 miRNA families were conserved in all the above three crop families. These conserved 14 miRNA families comprised only 31 members of miRNAs (Supplementary Table 1). In case of wheat, one miRNA (miR156f) was excluded because of high E-value during BLAST hit in Ensemble. Two miRNAs (169b and 169c) does not possess any repeat motif and 5 miRNAs (miR156e, miR827, miR399, mir167a and miR167b) had less than seven repeat motifs. Rest of the sequences generated a wide range of repeats which varied from a maximum of tetra-nucleotide repeat motif (NNNN) repeated 6 times (NNNN)6, to a minimum of di-nucleotide (NN) repeat motif repeated 27 times (NN)27. Similar strategy has been followed in Arabidopsis where miR159a, miR159b, miR166b, miR168, miR169c, miR169d, miR319, miR398, and miR167b were excluded due to high E-value and miRNA169b did not have any repeat motif and three miRNAs (miR827, miR393a, and miR399) possessed less than seven repeat motifs. Similarly, in rice, two miRNAs (miR827 and miR167b) were excluded due to high E-value and 3 miRNAs (miR827, miR393a and miR399) had less than seven repeat motifs. Similar to wheat miRNAs, 169b and 169c have no repeat motif in rice as well. However, overall, di-nucleotide repeats were found in high frequency (48), followed by tri-nucleotide (9), tetra nucleotide (5), penta- and hexa-nucleotide (4) repeats (Supplementary Table 1).

For validation, spanning to 31 nucleotides, we finally selected 13 sequences that had motif length equal/more than 12 and have been targeted for designing SSR primer pairs. The primer pairs of 13 miR-SSRs were tried on a panel of 37 wheat genotypes comprising of both heat tolerant and heat susceptible genotypes. All the 13 miR-SSRs designed amplified expected, clear, reproducible banding pattern in all the 37 wheat genotypes.

### Allelic diversity and molecular characterization of heat tolerant/heat susceptible wheat genotypes

The knowledge of nature and extent of genetic diversity/allelic diversity available in the germplasm helps the plant breeders for planning purposeful breeding programmes. Molecular markers have proven useful for their role in assessment of genetic variation in germplasm collections, characterization and evaluation of genetic diversity in crop germplasm [37, 38, 39]. A variety of random markers have been used in the study of genetic diversity/allelic diversity in wheat crop but there is hardly any report available where miRNA derived SSR markers have been used in the study of genetic diversity, molecular characterization of wheat germplasm. Therefore, we have tried 13 mi-RNA derived SSR markers on 37 wheat genotypes comprising of both heat tolerant and heat susceptible genotypes. The analysis revealed that a total of 30 alleles were detected using these 13 SSR markers. However, out of these 13 SSRs, only seven were found to be polymorphic SSRs (62.50% polymorphism) while SSRs designed for HT-156a, HT-156b, HT- 166c, HT-167c, HT-171, and HT-398 amplified only one allele i.e., were monomorphic (Table 2). The lowest amplicon size (127 bp) amplified for HT-171 SSR, and the highest amplicon size (229 bp) amplified for HT-160a SSR (Table 2). The analysis of PIC indicated that out of the seven polymorphic markers, the highest PIC value (0.375) depicted by miR156e-SSR and the lowest (0.164) by miR169j-SSR. Markers HT-169j, HT-160a, and HT-160b (Figure 1) were found superior for the analysis of overall genetic diversity at the respective loci as these markers showed more length variation among different genotypes used in the present study and therefore, are considered to be the best markers to reveal true genetic diversity and for their use in wheat germplasm characterization (Table 2 for a summary).

**Fig. 1.**
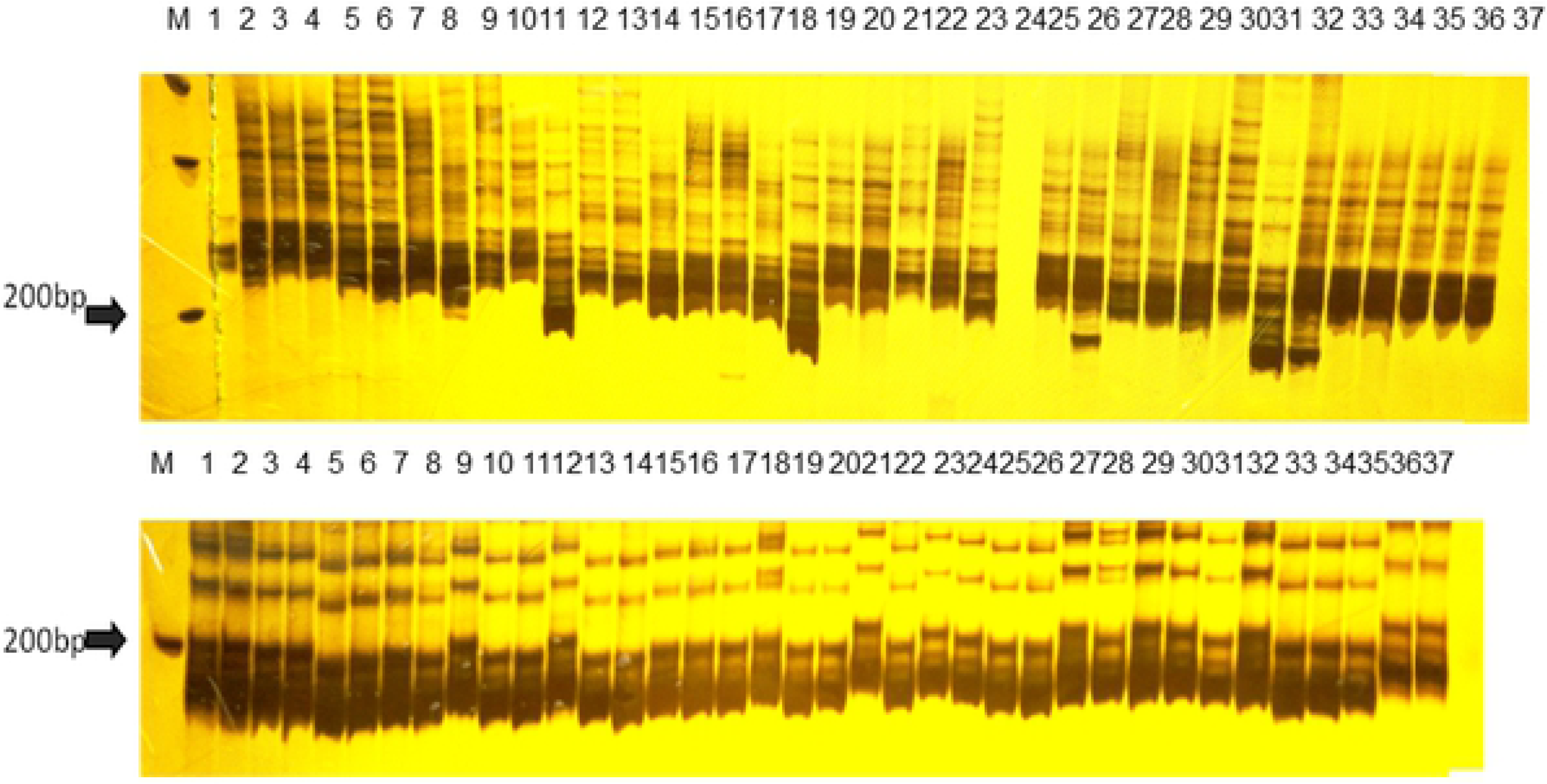
PAGE amplification profile of a set of 37 wheat genotypes showing length polymorphism for the SSR HT-160a and HT-160b. M-100 base pair ladder, 1-11; heat susceptible wheat genotypes panel, and 12-37; heat tolerant wheat genotypes panel.

**Table 2:**
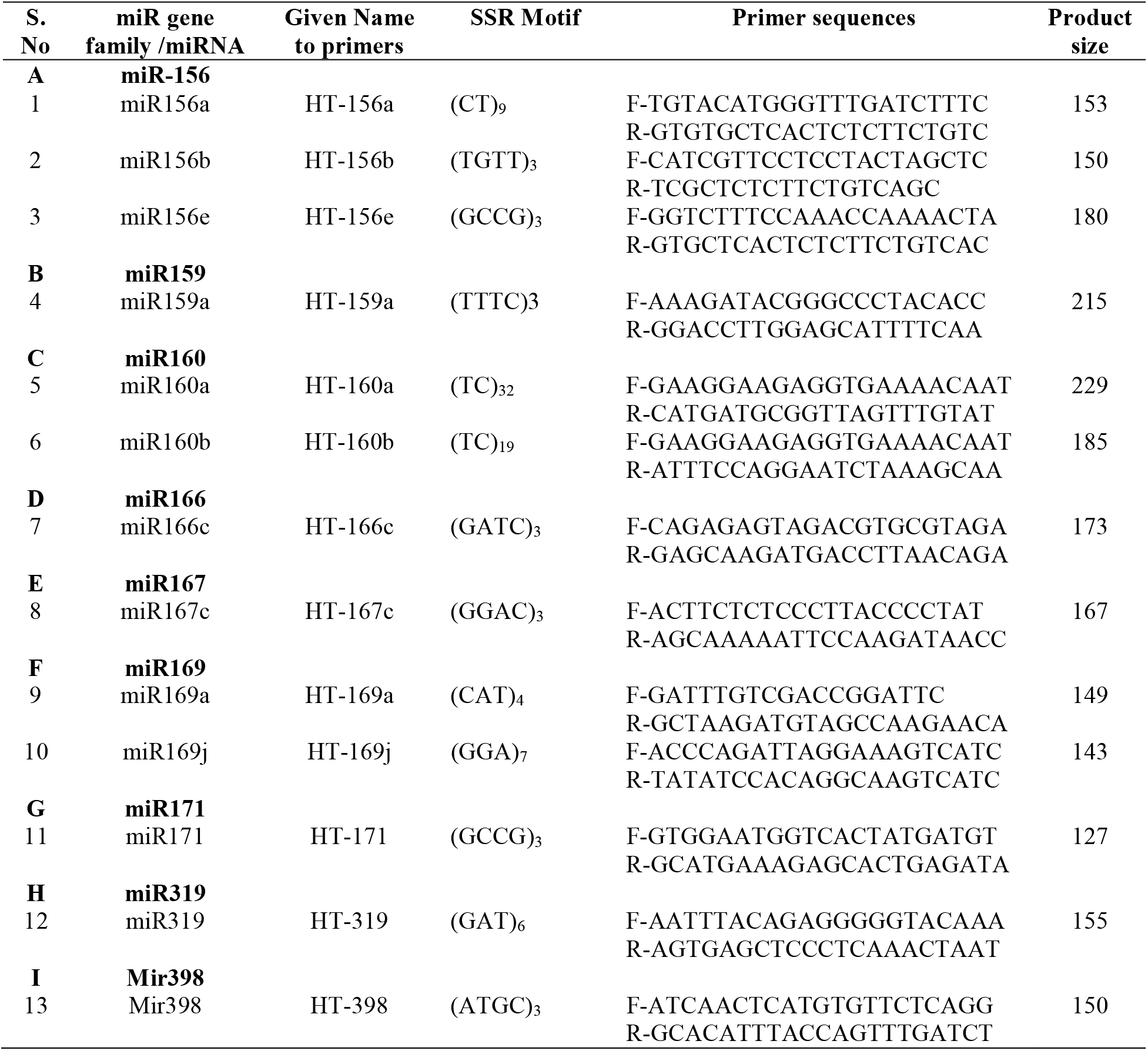
Details of 13 miRNA-SSRs including miR gene family with their miRNA, SSR Motif found in particular miRNA, primer sequences against identified miRNA–SSRs, and the product size at which particular primer was amplified.

### Population structure and genetic relationships

For the study of population genetic structure in a set of 37 wheat breeding lines/varieties, Bayesian clustering was executed using a software program “STRUCTURE” (Figure 2). Genotypic data of seven miRNA-SSRs on 37 wheat genotypes was analysed using this software. As the clustering model presumes the underlying existence of sub-populations “K clusters”, an Evano test was performed that predicted four sub-populations “K = 4” in our collection of 37 lines (Figure 2a). The analysis of four sub-populations identified by structural analysis indicated that 10, 12, 8, and 7 genotypes are present in four sub-populations “P1, P2, P3, and P4” respectively. Similar results were also obtained from PCoA by clustering the 37 wheat genotypes into four clear groups (Figure 2c). A significant genetic divergence was observed among the four sub-populations from each other (Table 3). A total of nine genotypes were found to be admixed.

**Fig. 2.**
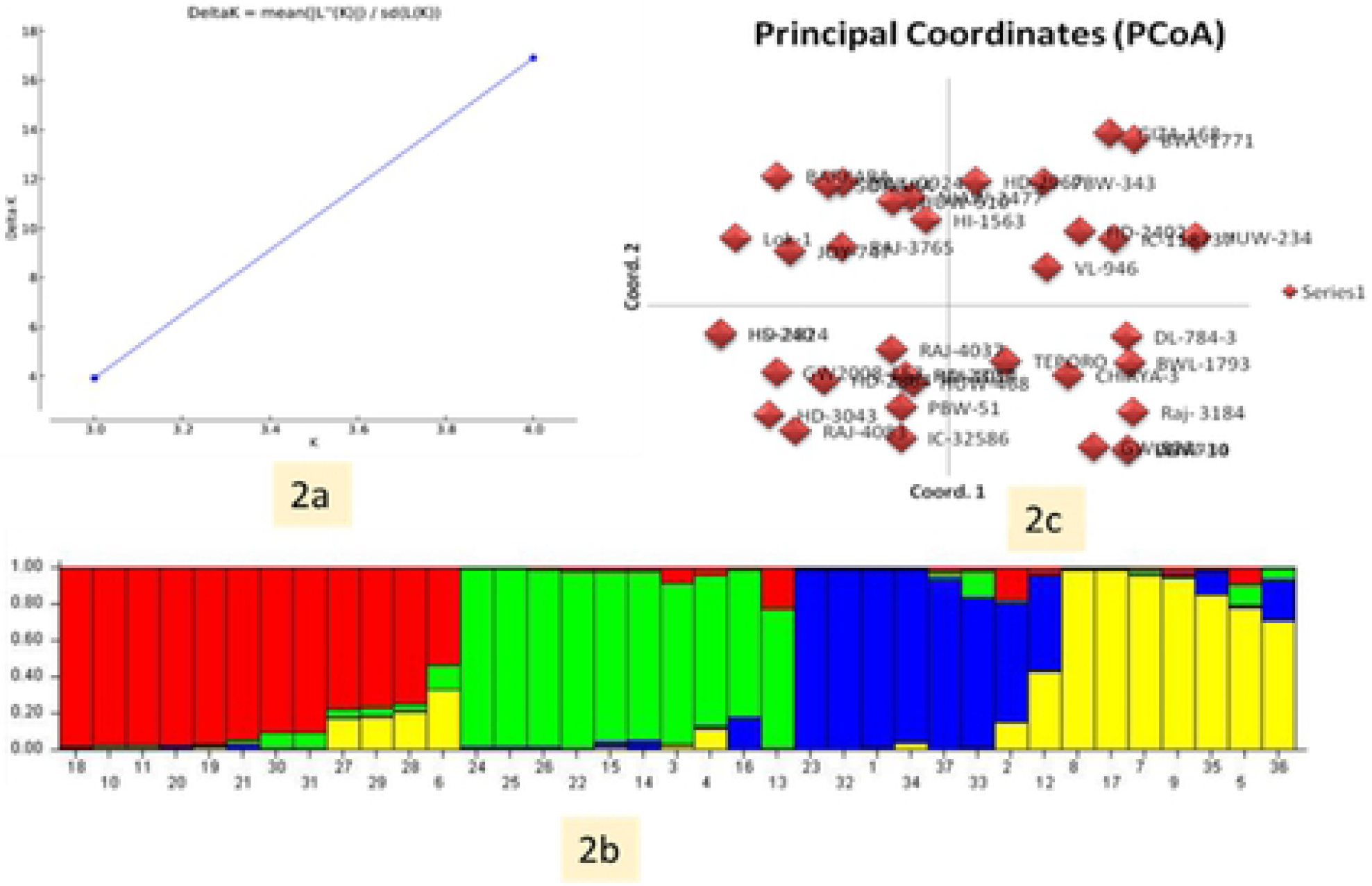
Estimation of the number of groups based on output from STRUCTURE-software. 2a: Δ K over K from 2–10 with the 13-miRNA-based SSR markers. 2b: Bar plot showing grouping of 37 genotypes into four different groups, and 2c: PCA results grouping of 37 wheat genotypes into four groups.

**Table 3:** Analysis of molecular variance using 13 miRNA-SSRs of the genetic differentiation among and within four subpopulations.

| Source | df | SS | MS | Est. Var. | % |
| --- | --- | --- | --- | --- | --- |
| Among Pops | 3 | 16.518 | 5.506 | 0.139 | 3% |
| Within Pops | 33 | 139.914 | 4.240 | 4.240 | 97% |
| Total | 36 | 156.432 |  | 4.379 | 100% |

### Genetic differentiation of populations

The information about four sub-populations identified by STRUCTURE analysis was used in GenAlex 6.5 software for AMOVA analysis and the genetic diversity indices. The AMOVA revealed that 3% of the total variation was found among/between sub-populations, while, the rest of variation (97%) was due to individuals/genotypes present within sub-populations (Table 3). These results demonstrated that genetic differentiation among sub-populations was low and within sub-populations was high. The mean values of different genetic parameters of the four subpopulations have been provided in Table 4.

**Table 4:** Mean of different genetic parameters including the number of different alleles (Na), the number of effective allele (Ne), Shannon’s index (I), diversity index (h), and unbiased diversity index (uh) in each subpopulation of the 37 genotypes.

| Pop | N | Na | Ne | I | H | uh | %P |
| --- | --- | --- | --- | --- | --- | --- | --- |
| Pop1 | 12.000 | 1.733 | 1.471 | 0.409 | 0.274 | 0.298 | 76.67 |
| Pop2 | 10.000 | 1.467 | 1.398 | 0.349 | 0.234 | 0.260 | 63.33 |
| Pop3 | 8.000 | 1.433 | 1.424 | 0.352 | 0.241 | 0.275 | 60.00 |
| Pop4 | 7.000 | 1.433 | 1.446 | 0.359 | 0.248 | 0.289 | 60.00 |
| Mean | 9.250 | 1.517 | 1.435 | 0.367 | 0.249 | 0.281 | 65.00 |

### Cluster analysis of 37 wheat genotypes

The data from miR-SSR profiling were also used to study the genetic diversity among the 37 genotypes. The results led to the clustering of 37 genotypes into four main groups/clusters (GI, GII, GIII and GIV) (Figure 3). There were 9, 7, 10, and 9 genotypes in group GI, GII, GIII, and GIV respectively. GI and GII included all heat stress tolerant genotypes, while group GIII and GIV contained a set of 6/4 and 5/4heat stress tolerant/ susceptible genotypes. This may be because of the same genetic background. For example, the parental genotypes PBW343 (heat susceptible) and its derived line Raj 4083 (heat tolerant) clustered together in GIII. Similarly, heat susceptible line HD2402 is a parental line of heat tolerant line Raj 3765 and clustered with it in GIV. The Jaccard’s similarity index between the pairs of wheat genotypes ranged from 32% to 89.5% with a mean similarity index of 55.9%. The principle coordinate analysis demonstrated that the 37 wheat genotypes can be separated distinctly from each other (Fig. 2c).

**Fig 3.**
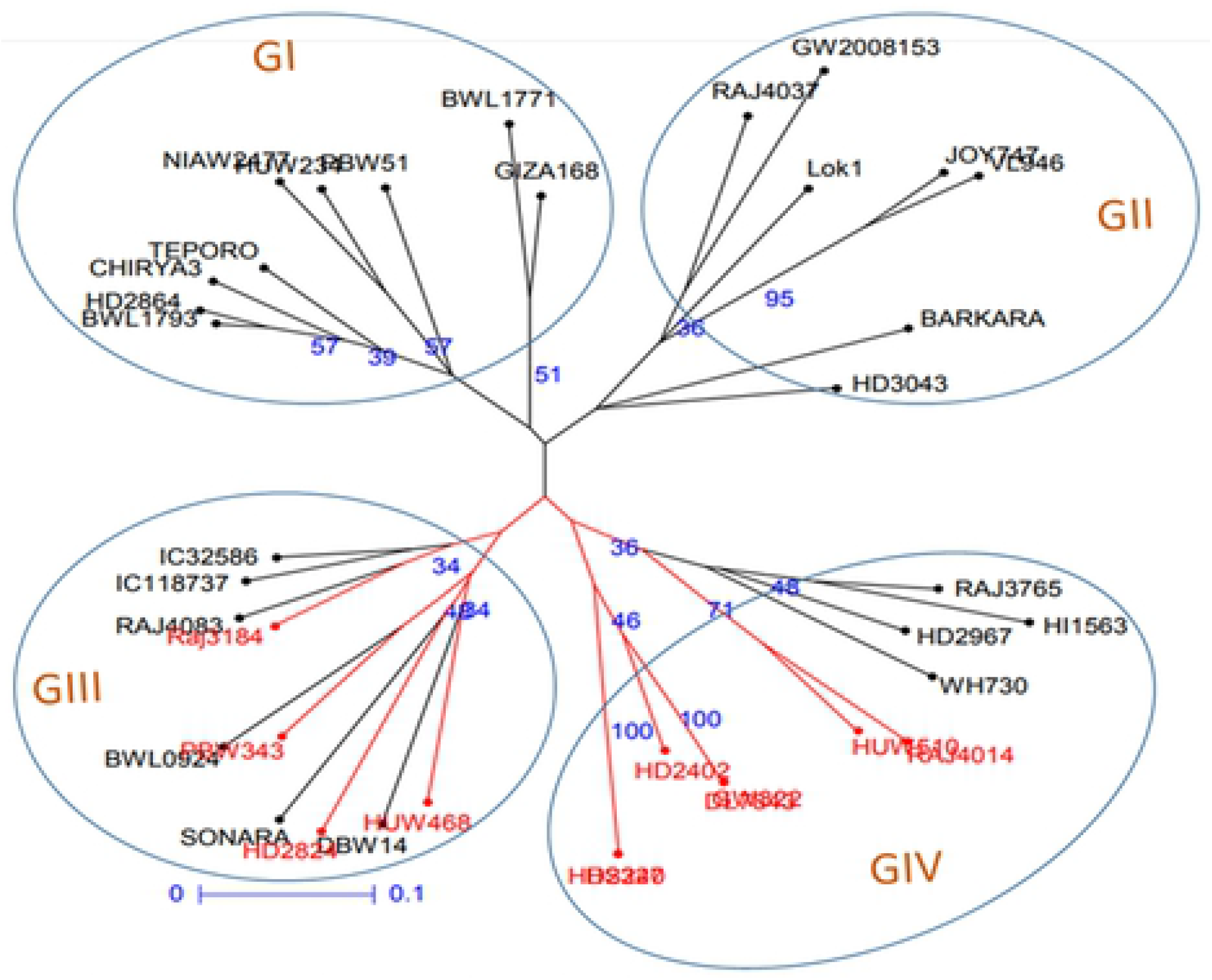
Unweighted neighbour-joining dendrogram showing grouping of 37 genotypes based on miRNA-SSR genotypic data.

## Discussion

miRNAs are considered very important small non-coding RNA molecules and can be differentiated from other class of RNAs by possessing characteristics like (i) miRNAs are derived from genomic loci distinct from other recognized genes, (ii) miRNAs are processed from transcripts, and (iii) miRNAs sequences are conserved in related organisms. Plant miRNAs have been described for the first time in *Arabidopsis thaliana and* later in other species. At present miRNAs have now been reported in dozens of plants species [40,19,3] and their sequences have been deposited in a publicly-available miRNA database, miRBase (http://www.sanger.ac.uk/cgi-bin/Rfam/mirna/browse.pl). Several miRNAs that have been identified and characterised in crop plants have been found involved in development, phyto-hormone signalling, flowering and sex determination and responses to biotic and abiotic stresses [40].

Lately, miRNA-based markers have been developed and used for various purposes in crop plants including study of genetic diversity [16]. Among a range of molecular markers, SSR markers have been widely used for genetic diversity studies, QTL mapping, GWAS and marker-assisted breeding (MAB) programs etc. Polymorphism in SSRs is due to change in the number of repeats motif/unit. SSRs present within the gene (genic SSRs) are more important for molecular breeding studies than random genomic SSRs. Most of them are from either protein coding regions or un-translated regions of plant genomes [41]. The occurrence of SSRs within the miRNAs and their role against different biotic and abiotic stress has been adopted as a key topic during last decade [42]. As widely known, miRNAs are a 19-24 nucleotides stretch of non-coding RNAs (ncRNAs), involved in regulation of gene expression in plants under different biotic and abiotic stresses [16, 43,42]. The position of SSRs within pre-miRNAs sequences is not fixed [44]. They also differ in their number and occurrence of repeats in pre-miRNAs in descendent order from mono-nucleotide to hexa-nucleotide repeats [44].

Genetic diversity analysis helps in characterization and conservation of genetic resource. SSRs within the miRNA precursors have been identified and used for the diversity analysis [16,45]. Heat stress, highly effecting world wheat crop production is an important abiotic stress. To over-come the diminishing effect there is a need to identify wheat varieties with a high tolerance level towards heat stress or improvement of cultivated wheat varieties through many approaches including both forward and reverse genetics. Recent literatures suggest a large set of different abiotic stress related miRNAs have been reported in many crops including miRNAs for heat stress in wheat [46]. In addition, a number of studies have been available with the characterization of heat stress tolerant wheat germplasm [47,48] with a coding region based SSR markers. Yet to the best of our knowledge, this is the first study of identification of heat responsive miRNA-SSRS and characterisation of a set of wheat genotypes using identified markers.

*In silico* studies have been conducted to identify miRNA-based SSRs in Arabidopsis [24] and rice [44,50,16] but it has not been explored in wheat. By taking into consideration the importance of miRNA-based SSRs in crop breeding, we screened heat responsive miRNA genes of wheat for the identification of SSR motifs and analysed the possibilities of difference in response of the germplasm to heat stress due to these SSRs. A number of QTLs have been identified on all seven homeologous groups/chromosomes in earlier mapping studies for heat stress in wheat [48]. Out of 13 designed miRNA-SSR primers selected from eight chromosomes of five homeologous group 3, 4, 5, 6 and 7 of wheat, seven were monomorphic and were not included for further analysis. It has been earlier reported that mutation rate is high in higher repeat number motifs resulting in the high level of polymorphism for those SSRs loci [24]. This change in polymorphism may be seen as an adaptive change in the differential gene expression. All the 37 wheat genotypes showed a multi-allelic pattern for the selected seven polymorphic SSR markers, which indicate their high level of genetic diversity in response to heat tolerance trait in wheat. From the dendrogram, GI and GII clustered contain all heat tolerant genotypes, however, GIII and GIV showed clustering of both heat tolerant and heat susceptible genotypes, which may due to the number of markers was not enough to cover the large genome of wheat or fewer genotypes from different geographical locations were used in this study. The differentiation of heat tolerant genotypes in cluster GI and GII indicated the usefulness and utility of micro-RNA SSR markers in genetic diversity studies in wheat and may indicate the candidate nature /functional nature of these micro-RNA sequences and hence SSRs for heat tolerance. These SSR may also show a strong association with heat tolerance using different methods of marker-trait association or regression methods.

As a result of this study, all seven miR-SSR markers were polymorphic but out of these only three were able to distinguish two contrasting groups of wheat genotypes for heat stress. Expansions or contractions of the repeat motifs at the SSRs have been reported to play a major role in different gene activities (ref?). Therefore, this variation in length may cause respective changes in gene expression patterns, increasing/decreasing the initial stress tolerance level i.e., phenotype change without gene alteration.

## Conclusion

In this study, we identified three miRNA based SSR markers that can differentiate contrasting set for heat tolerance in wheat. We conclude that the same miRNA for heat stress can be different in expression in different genotypes. The length of SSRs might be responsible for the variation in the expression, as allelic variation was more in susceptible genotypes than the tolerant genotypes. The number of motif repeats are responsible for their higher level of polymorphism. These miRNA-SSRs will play an important role in serving as a source of highly informative molecular markers and aids as a reference for marker assisted breeding in plants.

## Author’s contribution

Conceptualization and Data curation was done by ST. AK and TG help in methodology and genotyping, respectively. ST and RRM analysed the data and prepared the manuscript. RP and RRM revised the manuscript. All authors read and approved the manuscript.

## Acknowledgement

ST acknowledges receipt of funding support from Science and Engineering Research Board (SERB), New Delhi (Award Number: **PDF/2016/002667**).

